# H3K27me3-mediated silencing of Wilms Tumor 1 supports the proliferation of brain tumor cells harboring the H3.3K27M mutation

**DOI:** 10.1101/114082

**Authors:** Dong Fang, Haiyun Gan, Liang Cheng, Jeong-Heon Lee, Hui Zhou, Jann N. Sarkaria, David J. Daniels, Zhiguo Zhang

## Abstract

The lysine 27 to methionine mutation of histone H3.3 (H3.3K27M) is detected in over 75% of diffuse intrinsic pontine glioma (DIPG). The H3.3K27M mutant proteins inhibit H3K27 methyltransferase complex PRC2, resulting in a global reduction of tri-methylation of H3K27 (H3K27me3). Paradoxically, high levels of H3K27me3 were also detected at hundreds of genomic loci. However, it is not known how and why H3K27me3 is redistributed in DIPG cells. Here we show that lower levels of H3.3K27M mutant proteins at some genomic loci contribute to the retention of H3K27me3 peaks. But more importantly, Jarid2, a PRC2-associated protein, strongly correlates the presence of H3K27me3 and relieves the H3.3K27M-mediated inhibition *in vivo and in vitro.* Furthermore, we show that H3K27me3-mediated silencing of tumor suppressor gene Wilms Tumor 1 (WT1) supports the proliferation of DIPG cells and reaction of WT1 inhibits DIPG proliferation. Together, these studies reveal mechanisms whereby H3K27me3 is retained in the environment of global loss of this mark, and how persistence of this mark contributes to DIPG tumorigenesis.

## Introduction

Diffuse intrinsic pontine glioma (DIPG) is the most aggressive primary malignant brain tumor found in children (Louis et al., 2007). The median survival time after diagnosis is approximately one year with no cure in sight (Dellaretti et al., 2012). Recent studies have identified, in about 75% of DIPG cases, a somatic mutation of the *H3F3A* gene, leading to a lysine 27 to methionine mutation at histone H3 variant H3.3 (H3.3K27M) (Schwartzentruber et al., 2012; Sturm et al., 2012; Wu et al., 2012, 2014). In addition, *HIST1H3B* or *HIST1H3C,* one of the 13 genes encoding canonical histone H3.1/H3.2, is also mutated with the same K to M changes in a small fraction of DIPG tumors (Castel et al., 2015; Fontebasso et al., 2014; Solomon et al., 2016). Introducing the mutation in human or mice can induce tumor formation, indicating that it is a driver of tumorigenesis. Therefore, it is critically important to understand how the histone H3K27M mutation drives tumorigenesis.

H3.3 is a histone H3 variant that differs from H3.1/H3.2 by only four or five amino acid residues (Campos and Reinberg, 2009; Maze et al., 2014; Szenker et al., 2011). Compared to canonical histone H3.1/H3.2 that are assembled into nucleosomes in a replication-coupled process, histone variant H3.3 is assembled into nucleosomes in a replication independent manner (Ahmad and Henikoff, 2002a; Tagami et al., 2004). H3.3 plays important roles in a variety of cellular processes. For instance, H3.3 is involved in the epigenetic memory of actively transcribed genes in nuclear transferred *Xenopus* nuclei (Ng and Gurdon, 2008). Paradoxically, H3.3 is also important for heterochromatin formation during mouse development (Banaszynski et al., 2013; Jang et al., 2015; Lin et al., 2013) and during oncogene induced senescence (Duarte et al., 2014; Lee and Zhang, 2016; Rai et al., 2014; Zhang et al., 2007), as well as for silencing of endogenous retroviral elements in mouse embryonic stem cells (Elsässer et al., 2015). H3.3 at gene bodies is enriched at actively transcribed genes compared to lowly expressed genes (Ahmad and Henikoff, 2002b; Chen et al., 2013; Deaton et al., 2016), supporting a role of H3.3 in gene transcription. H3.3 has also been shown to play a role in DNA replication during replication stress (Adam et al., 2013; Frey et al., 2014). It is proposed that the context-dependent role of H3.3 in different cellular processes are likely linked to its assembly at distinct chromatin regions by different histone chaperones and regulators (Burgess and Zhang, 2013).

Like other histone H3 proteins, H3.3 is modified post-transcriptionally via the same enzymes, as very few H3.1 and H3.3 specific enzymes have been reported (Guo et al., 2014; Jacob et al., 2014; Wen et al., 2014). These modifications, including acetylation and methylation, are associated with and/or regulate different cellular processes (Jenuwein and Allis, 2001; Klose and Zhang, 2007; Pokholok et al., 2005; Tan et al., 2011; Turner, 2000). H3K27 can be acetylated (H3K27ac) by p300/CBP (Suka et al., 2001) and this modification marks active enhancers in combination with H3K4 mono-methylation (H3K4me1) (Calo and Wysocka, 2013; Creyghton et al., 2010; Heintzman et al., 2009; Shen et al., 2012). H3K27 can also be mono-, di-and trimethylated (H3K27me1/me2/me3). H3K27me2/me3, catalyzed by the Polycomb repressive complex2 (PRC2) (Pirrotta, 1998), is enriched at promoters of silenced genes and plays an important role in the silencing of developmentally regulated genes during cell differentiation (Banaszynski et al., 2013; Cao et al., 2002; Margueron and Reinberg, 2011; Simon and Kingston, 2013).

The H3K27 methyltransferase complex PRC2, which consists of four core subunits, Ezh2, Suz12, EED, and RbAP46/48, methylates nucleosomal histone H3K27. In *Drosophila* cells, the PRC2 complex is recruited to chromatin through DNA binding proteins to Polycomb response elements (PRE), which contain specific *cis*-regulatory DNA sequences (Simon et al., 1993). In human cells, however, consistent PRE elements have not been identified and the core subunits of PRC2 complex do not have DNA sequence specific binding abilities, suggesting that human PRC2 is recruited in a more complex manner than *Drosophila* PRC2 (Mendenhall et al., 2010; Woo et al., 2010). Indeed, several mechanisms including protein binding partners, non-coding RNAs as well as coding RNAs have been proposed for the recruitment of PRC2 to specific loci for H3K27 methylation (Kaneko et al., 2013; Kanhere et al., 2010; Khalil et al., 2009; Zhao et al., 2010). For instance, PRC2 forms a complex with Jarid2, which targets PRC2 to specific loci (Peng et al., 2009; Shen et al., 2009). Jarid2, a member of Jumonji (Jmj) family, contains a JmjN domain, a JmjC domain, a zinc finger domain, an RNA-binding region (RBR) and a DNA-binding domain, also called AT-rich interaction domain (ARID). Unlike other members of the Jmj family that catalyze the demethylation of lysine residues, Jarid2 does not contain the conserved residues required for iron and ascorbate binding that are essential for the demethylation activity (Pasini et al., 2010; Takeuchi et al., 2006). Jarid2 is enriched at CGG- and GA-containing sequences (Peng et al., 2009), and also binds to long non-coding RNA (lncRNA) through RBR *in vivo* and *in vitro* (Kaneko et al., 2014). For instance, Jarid2 interacts with MEG3 (Rocha et al., 2008), an lncRNA encoded by the imprinted DLK1-DIO3 locus (Kaneko et al., 2013). It is likely that Jarid2 targets PRC2 to specific loci through its DNA as well as RNA binding activities. Jarid2 is highly expressed in embryonic stem (ES) cells and is required for mouse embryogenesis, including neural tube formation (Takeuchi et al., 1995) and heart development (Mysliwiec et al., 2011, 2012). *In vitro,* Jarid2 enhances the enzymatic activity of PRC2 complex (Sanulli et al., 2015; Son et al., 2013). As a partner of PRC2, Jarid2 facilitates the recruitment of PRC2 to its target genes, stimulates the H3K27 methyltransferase activity, and plays a role in silencing of key regulatory genes during mouse ES cell differentiation.

We and others have found that the levels of H3K27me3 are reduced dramatically in DIPG cells as well as in any tested cell type or organisms expressing H3.3K27M mutant proteins, including mice and *Drosophila* (Chan et al., 2013a, 2013b;Herz et al., 2014; Lewis et al., 2013). Several lines of evidence support the idea that the global reduction of H3K27me3 in DIPG cells is due to inhibition of PRC2 by H3.3K27M mutant proteins. *In vivo,* Ezh2 was enriched at H3.3K27M mono-nucleosomes isolated from cells (Bender et al., 2013; Chan et al., 2013b). *In vitro,* PRC2 complex binds to H3.3K27M peptides with high affinity (Justin et al., 2016; Lewis et al., 2013), and H3.3K27M peptides or mononucleosomes inhibit the enzymatic activity of PRC2 complex (Bender et al., 2013; Jayaram et al., 2016; Jiao and Liu, 2015; Justin et al., 2016). The crystal structure of H3K27M peptide-PRC2 complex reveals that the H3K27M peptide binds to the active site of Ezh2 in a canonical manner, with the side chain of methionine positioned in the lysine access channel, which provides mechanistic explanation for the high binding affinity of PRC2 by H3.3K27M mononucleosomes (Jiao and Liu, 2015; Justin et al., 2016). In addition to the global loss of H3K27me3, we and others also observed that H3K27me3 peaks were detected at the promoters of hundreds of genes in DIPG xenograft cell lines (Bender et al., 2013; Chan et al., 2013b). Strikingly, gene etiology analysis indicates that these genes associated with H3K27me3 in DIPG cells are enriched in cancer pathways (Chan et al., 2013b). However, it is largely unknown how H3K37me3 are present at these specific genes in the environment of a global loss of this mark. Moreover, the functional implications for the presence of H3K27me3 in tumorigenesis are also unexplored.

Here, we show that locus-specific retention of H3K27me3 in DIPG cells is a conserved phenomenon detected in two DIPG xenograft cell lines and one primary tumor sample. Moreover, we provide two non-exclusive mechanisms for the locus-specific-retention of H3K27me3 in the environment of global loss in DIPG cells. First, H3.3K27M mutant proteins are low at these gene promoters. Second, Jarid2, a partner for the PRC2 complex, can relieve the inhibitory effect of H3.3K27M mutant proteins. Functionally, we show that H3K27me3 mediated silencing of tumor suppressor gene Wilms Tumor 1 (WT1) is required for the proliferation of DIPG cells.

## Results

### H3K27me3 was detected in DIPG patient tissue and cell lines with H3.3K27M mutation

In addition to dramatic global reduction in H3K27 methylation as detected by Western blot and ChIP-seq assays, we also paradoxically observed the enrichment of H3K27me3 within the promoters of hundreds of genes in DIPG cell line (SF7761) harboring the H3.3K27M mutation (Chan et al., 2013b). To determine whether the genes associated with H3K27me3 in SF7761 are also methylated in other DIPG tumor line and in primary DIPG tissue, we performed chromatin immunoprecipitation coupled with next-generation sequencing (ChIP-seq) in another DIPG xenograft cell line (SF8628), and one pontine glioma tissue sample with H3.3K27M mutation (Figure S1A). We identified 2,356 H3K27me3 ChIP-seq peaks in SF7761, 5,080 in SF8628 and 12,033 peaks in DIPG pontine tissue. When normalized against spike-in chromatin, the majority of these peaks exhibited reduced H3K27me3 levels. However, about 9% of H3K27me3 peaks in SF7761 and SF8682 and 6% of H3K27me3 in primary tissue exhibited similar or even higher levels of H3K27me3 peaks compared to their reference counterpart (Figure S1B). These results suggest that H3K27me3 peaks detected in DIPG cells are likely due to both the retention of H3K27me3 peaks in tumor-initiating cells as well as “gain” of H3K27me3 in tumor cells. In the rest of text, we did not address these two possibilities. Instead, we focus on analysis the mechanisms for the presence of H3K27me3 peaks and the functional consequence of these peaks for proliferation of DIPG cells.

Of all these H3K27me3 ChIP-seq peaks in two DIPG lines and one DIPG tissues, 1,088 H3K27me3 ChIP-seq peaks in SF7761, 2,729 in SF86828 and 3,431 in primary pontine tissue were localized at promoters. Importantly, 676 H3K27me3 peaks at/close to promoters overlapped among these three samples (Figure 1A-1B). Of these 676 peaks, H3K27me3 surrounding the transcription start site (TSS) exhibited a similar pattern among the DIPG cell lines and patient tissue (Figure 1C and 1D). These findings indicate that a large fraction of H3K27me3 peaks are likely present in all DIPG samples with the H3.3K27M mutation.

**Figure 1.**
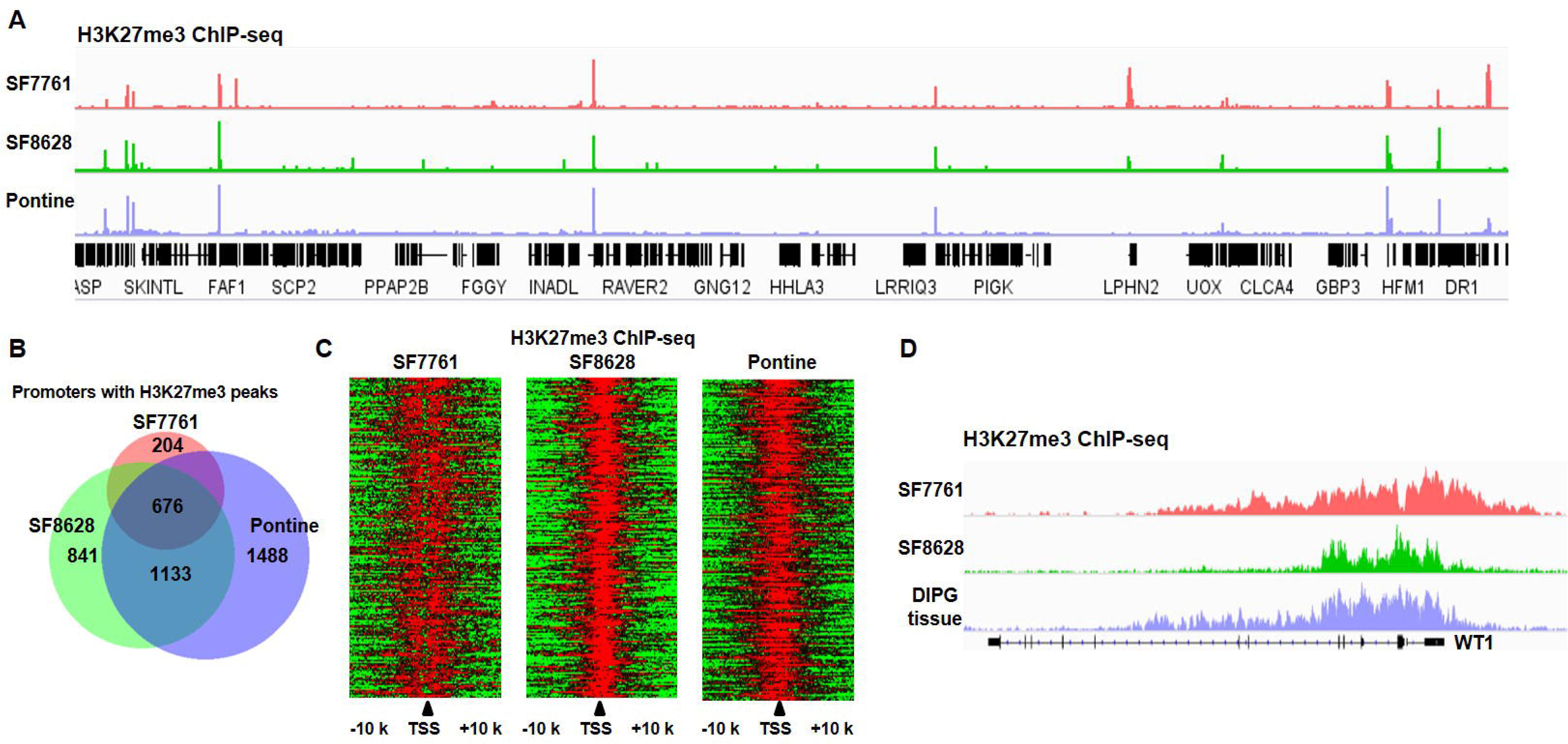
H3K27me3 peaks are present in both DIPG cell lines and tissue. (**A**) Representative Integrative Genomics Viewer shows the distribution of H3K27me3 in two DIPG cancer cell lines (SF7761 and SF8628) and one DIPG primary tissue. RefSeq genes are shown at the bottom. (**B**) Venn diagram illustration represents the promoters with H3K27me3 peaks among SF7761, SF8628, and pontine DIPG tissue. (**C**) Heatmaps represent the signal of H3K27me3 from 10Kb upto 10Kb down-stream of TSS of 676 genes (B) that have H3K27me3 at the promoters in SF7761, SF8628, and one DIPG tissue. (**D**) Genome browser track examples for the occupancy profiles of H3K27me3 at Wilms Tumor 1 (WT1) locus.

### H3.3K27M mutant proteins are enriched at highly expressed genes and exhibit an inverse relationship with H3K27me3

The global loss of H3K27me3 in DIPG cells is linked to the inhibition of PCR2 by H3.3K27M mutant proteins. However, it was not known whether H3.3K27M mutant proteins have any role in the retention/gain of H3K27me3 in DIPG cells. We analyzed the localization of H3.3K27M mutant proteins using H3K27M specific antibody (Figure S2A). The H3.3K27M mutant proteins were enriched at actively transcribed genes compared to lowly expressed genes in both SF7761 and SF8628 cells, a pattern that is consistent with the localization of wild type H3.3 proteins on chromatin detected in other cell lines (Figure 2A-B). Under the same conditions, ChIP-seq signals of H3.3K27M mutant proteins were not detected in human neural stem cells (NSCs) (Figure 2C). These results suggest that H3.3K27M ChIP-seq peaks were specific and that H3.3K27M mutant proteins are assembled into nucleosomes normally. Next, we compared the levels of H3.3K27M mutant proteins at gene promoters with high and low H3K27me3. Using human NSC and neural progenitor cells (NPCs) as reference, we identified 9,305 promoters associated with H3K27me3 in either reference cells or DIPG tumor lines. We observed that the levels of H3.3K27M mutant proteins were lower at the 2000 gene promoters associated with high H3K27me3 than the ones at 2000 gene promoter genes with low levels of H3K27me3 in both SF7761 and SF8628 (Figure 2D-G). These results suggest that H3.3K27M mutant proteins are likely assembled into silent chromatin regions less efficiently and thereby help retain H3K27me3 at these regions.

**Figure 2.**
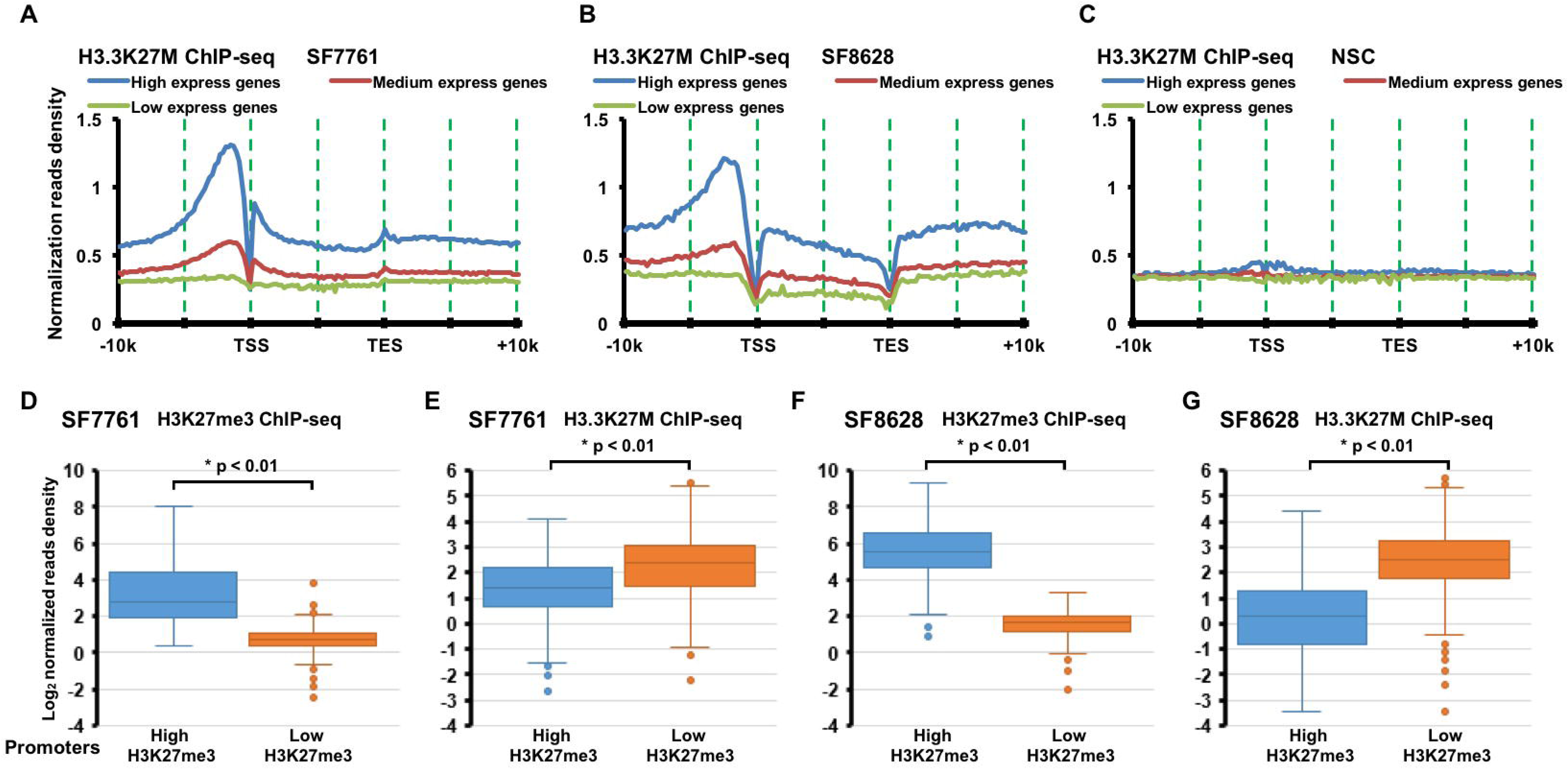
H3.3K27M levels on chromatin are inversely correlated with levels of H3K27me3 in both SF7761 and SF8628. (**A - C**) H3.3K27M mutant proteins are enriched at highly transcribed genes compared to lowly-expressed genes in DIPG cells. The average read density of H3.3K27M ChIP-seq in SF7761 (A), SF8628 (B), and NSC (C) from 10Kb upstream of TSS to 10Kb downstream of TES is calculated. The read density was normalized to RPKM (Reads Per Kilo-base per Million mapped reads). The entire human genes were split into three groups according to their expression levels in the corresponding cell lines: highest expressed genes, medium expressed genes, and low expressed genes. (**D - G**) The levels of H3K27me3 and H3.3K27M show an inverse-relationship. 9,305 promoters with H3K27me3 ChIP-seq peaks were identified in either of NSC (6845), NPC (5007), SF7761 (1088) or SF8628 (2729). The top 2000 promoters with highest H3K27me3 and 2000 promoters with the lowest H3K27me3 were chosen from these 9305 promoters to calculate H3K27me3 read density in SF7761 (D) and SF8628 cells (F). In addition, H3.3K27M ChIP-seq read density in SF7761 (E) and SF8628 cells (G) were also calculated. P value was calculated by two-tailed Student’s t test.

We also observed that 367 and 257 H3K27me3 peaks were in the vicinity of H3.3K27M mutant proteins in SF7761 and SF8628 cells, respectively (Figure S2B-C, data not shown). When aligned at the peak center, H3K27me3 at the H3.3K27M centers were significantly lower than the surrounding regions (Figure S2B-S2G). These results are consistent with the idea that the global loss of H3K27me3 is due to the inhibition by H3.3K27M mutant proteins. These results also suggest that other mechanism may exist to overcome the inhibition of H3.3K27M mutant proteins at these regions.

### Depletion of Jarid2 results in a reduction of the residual H3K27me3 in DIPG cells

We made a serendipitous observation suggesting that Jarid2 is responsible for the presence of H3K27me3 peaks in cells expressing H3.3K27M mutant proteins, when we analyzed H3K27me3 levels in mES cells knocked in with a Flag-tagged H3.3K27M mutation (data not shown). To determine whether Jarid2 and Ezh2 are required for residual H3K27me3 in DIPG cells, we depleted Ezh2 and Jarid2 in two DIPG cell lines and observed a reduction of residual H3K27me3 by Western blot (Figure 3A). The reduction of H3K27me3 was also validated at two loci with Jarid2 and H3K27me3 through ChIP-PCR (Figure 3B and S3A). Interestingly, we observed that depletion of Jarid2 led to reduction of Ezh2 at these two loci (Figure 3C and S3B), whereas deletion of Ezh2 had no apparent effect on the binding of Jarid2 (Figure 3D and S3C), consistent with the idea that Jarid2 is involved in recruiting Ezh2 to these loci for H3K27me3.

**Figure 3.**
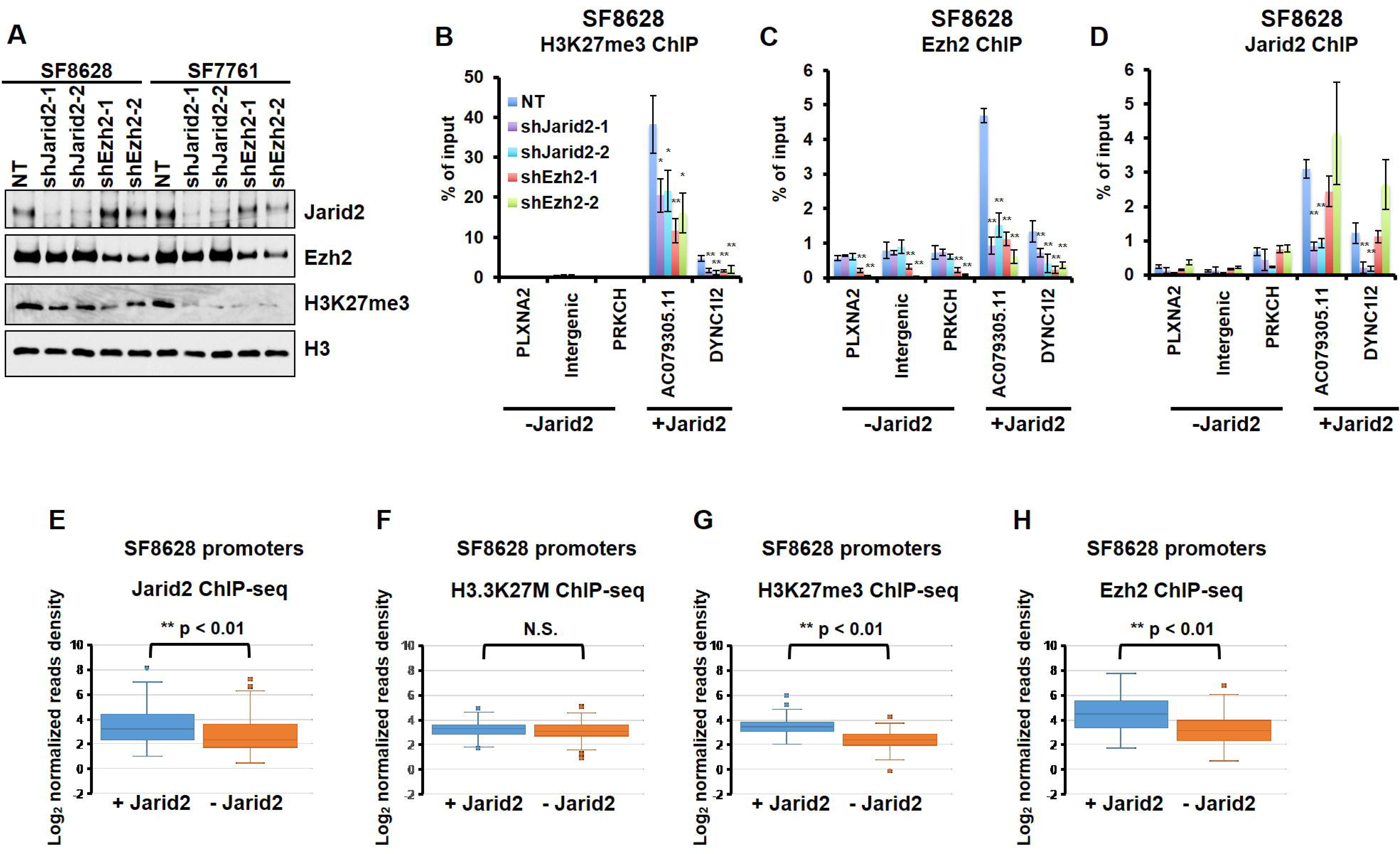
Jarid2 contributes to the presence of H3K27me3 in cells with H3.3K27M mutation. (**A - D**) Depletion of Ezh2 or Jarid2 leads to reduction of H3K27me3 levels in DIPG cells with H3.3K27M mutation. Ezh2 or Jarid2 was depleted in SF7761 or SF8628 cells. The total levels of H3K27me3, Jarid2 and Ezh2 was analyzed by Western blot (A). The levels of H3K27me3 (B), Ezh2 (C) and Jarid2 (D) at specific loci were analyzed by ChIP-PCR. AC079305.11 and DYNC1I2 are loci where H3K27me3, Ezh2 and Jarid2 are enriched. At Intergenic region, PLXNA2 and PRKCH, H3K27me3, Ezh2 and Jarid2 were low based on analysis of ChIP-seq and were used as controls. Data are mean ± SD (N = 3 independent replicates, ^*^P < 0.05, ^**^P < 0.01). NT, Non-Targeted control. (**E - H**) H3K27me3 levels are positively correlated with Jarid2 in the presence of H3.3K27M mutant proteins. All gene promoters (4195) associated with H3K27me3 in both reference cells (NSC and NPC) and SF8628 were first combined as reference H3K27me3 peaks in DIPG tumor initiating cells, of which 331 gene promoters have H3.3K27M peaks in SF8628 cells (see Figure S3D). These 331 genes were sub-grouped based on Jarid2 ChIP-seq peaks in SF8628 cells, and Log_2_ normalized reads density for Jarid2 (E), H3.3K27M (F), H3K27me3 (G), and Ezh2 (H) at the promoters with (+Jarid2) or without (- Jarid) Jarid2 ChIP-seq peaks were calculated and shown as Boxplot. P value was calculated by two-tailed Student’s t test. N.S. Not Significant.

### Jarid2 is enriched at gene promoters with H3K27me3 in DIPG cells

Next, we tested whether Jarid2 is enriched at the promoters containing both H3K27me3 and H3.3K27M mutant proteins. To do this, we performed Ezh2 and Jarid2 ChIP-seq in SF7761 and SF8628 cells, but analyzed these ChIP-seq results in each cell line independently. Similar results/conclusions were obtained from each line (see Figure 3E-H and Figure S3E-F). To analyze the relationships among the levels of Jarid2, Ezh2, H3.3K27M and H3K27me3 in SF8682 cells, we started to obtain the H3K27me3 peaks in “tumor initiating cells” or reference cells before the loss of H3K27me3 due to the expression of H3.3K27M mutant proteins. Because the expression of H3.3K27M in human neural progenitor cells resulted in neoplastic transformation and mouse neural stem cells are proposed to be the cell origin of DIPG (Funato et al., 2014), we chose human NSCs and NPCs as reference. Briefly, we first identified all gene promoters with H3K27me3 in reference cells (NSC and NPC) or SF8628 cells (Figure S3D). These 4,195 H3K27me3 peaks were assumed to be present in tumor initiating cells” of SF8682 tumor before the introduction of H3.3K27M mutant proteins. Of these 4195 peaks, 331 gene promoters contained both H3K27me3 peaks in the “tumor initiating cells” and H3.3K27M ChIP-seq peaks in SF8628 cells. Of these 331 peaks, we observed that 167 peaks overlapped with Jarid2 ChIP-seq peaks in SF8628 cells. We then compared H3.3K27M, Jarid2, H3K27me3, and Ezh2 ChIP-seq reads densities among the 331 peaks with (167) and without (164) Jarid2 ChIP-seq peaks in SF8682 cells (Figure 3E). We observed that in SF8628 cells, the H3.3K27M ChIP-seq reads density was similar between genes promoters associated and not associated Jarid2 (Figure 3F). In contrast, the H3K27me3 (Figure 3G) and Ezh2 (Figure 3H) were significantly higher at gene promoters with Jarid2 peaks than the ones without Jarid2 peaks. Similar results were obtained when we analyzed the relationships among Jarid2, Ezh2, H3.3K27M and H3K27me3 levels in SF7761 cells (Figure S3E and S3F). These results strongly suggest that the presence of Jarid2 can relieve the inhibition of PRC2 by H3.3K27M mutant proteins.

### H3.3K27M peptide inhibits the activity of PRC2-Jarid2 complex less efficiently

To test whether Jarid2 can relieve the inhibitory effect of H3.3K27M on PRC2, we first compared the inhibitory effect of the H3K27M peptide on the enzymatic activity of recombinant PRC2 core complex in the presence or absence of Jarid2. As reported (Lewis et al., 2013), the H3K27M peptide inhibited the enzymatic activity of core PRC2 complex towards mononucleosomes (IC50 = 73.2 ± 3.9 μM). Moreover, Jarid2 enhanced the enzymatic activity of the core PRC2 complex. Importantly, the H3.3K27M peptide exhibited a lesser inhibitory effect on the PRC2-Jarid2, with a 4.4 fold increase in IC50 compared to core PRC2 complex (IC50 = 256 ± 13.3 μM) (Figure 4A - 4C). The corresponding H3 wild type peptide showed no obvious effects on the enzymatic activity of PRC2 with or without Jarid2. These results show that H3.3K27M mutant proteins inhibit the enzymatic activity of the Jarid2-PRC2 complex less efficiently.

**Figure 4.**
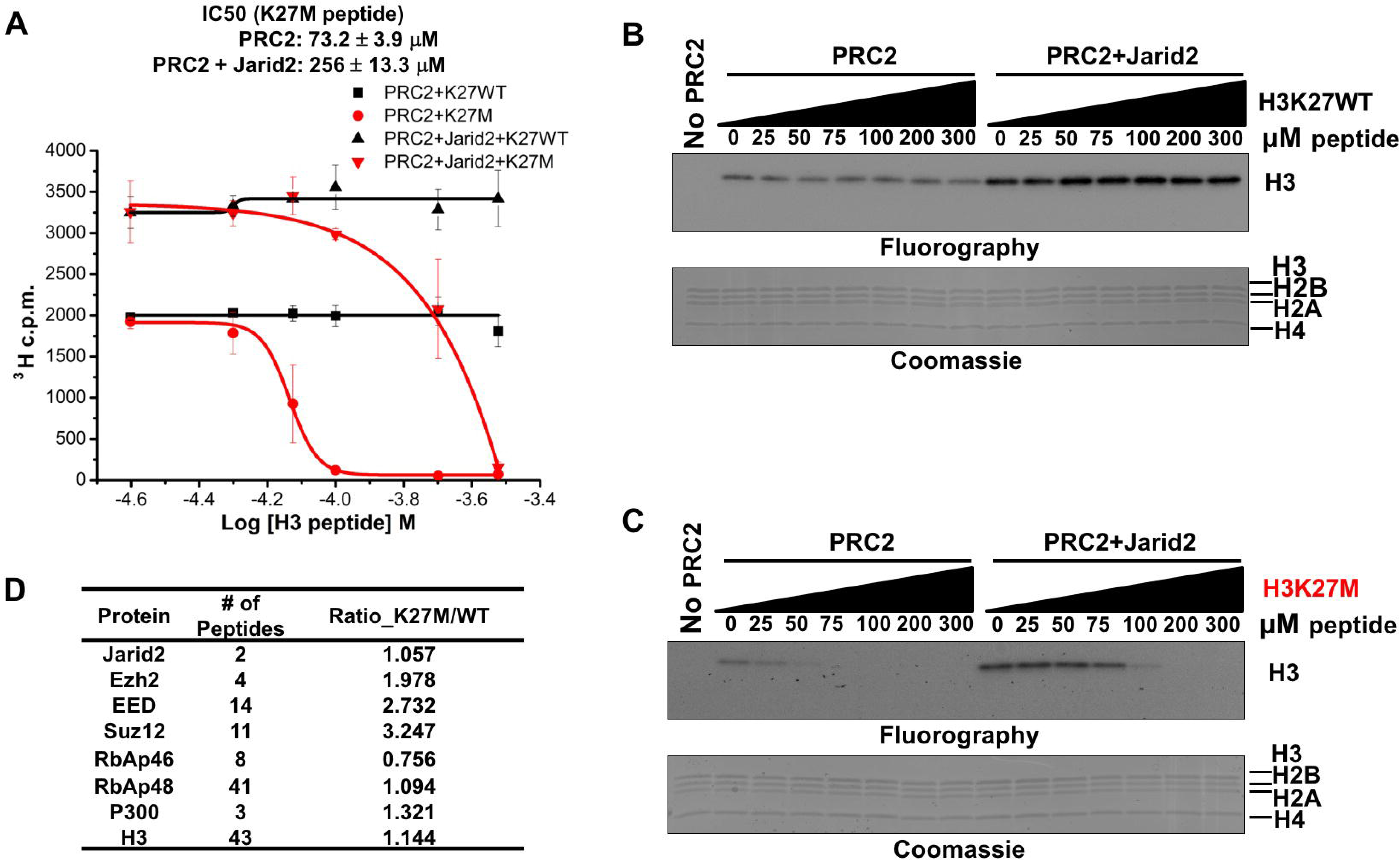
Jarid2 relieves the H3.3K27M-mediated inhibition of the enzymatic activity of PRC2 core complex. (**A**) Enzymatic activities of the H3K27 methyltransferase PRC2 with or without Jarid2 were measured using histone H3-containing recombinant mononucleosomes in the presence of increasing amounts of H3K27M peptides or its corresponding H3 wild type peptides. Data are mean ± SD (N = 3 independent replicates). c.p.m., counts per minute. (**B and C**) Autoradigraphy of the methyltransferase reactions with wild type H3.3K27 peptide (B) and H3.3K27M peptide (C) shows the H3K27M peptide inhibited Jarid2-PRC2 less efficiently. The in vitro methyltransferase assays were performed as described in (A). (**D**) Ezh2, EED and Suz13, but not Jarid2 were enriched on the H3.3K27M-containing mononucleosomes. HEK293T cells expressing Flag-tagged-H3.3 WT or H3.3K27M mutant proteins were grown in light-or heavy-isotope-labeled medium, respectively, and were mixed equally. Mononucleosomes were immunoprecipitated (IP) from mixed cells using antibodies against the FLAG epitope and co-IPed proteins were analyzed by LC-MS/MS. The SILAC ratio was the fold-enrichment of proteins in H3.3K27M-mononucleosome IP over WT H3.3-mononuclesome IP.

One possibility for the reduced inhibition of Jarid2-PRC2 complex by H3.3K27M mutant proteins is because the Jarid2-PRC2 complex binds to H3.3K27M-containing nucleosomes less efficiently. To test this idea, we performed mononucleosome immunoprecipitation and analyzed the relative abundance of PRC2 core complex and Jarid2 that co-purified with H3.3K27M- and wild type H3.3-containing mononucleosomes using SILAC coupled to LC-MS/MS. We observed that Ezh2, EED, and Suz12 were similarly enriched on the H3.3K27M-containing mononucleosomes compared to wild type H3.3 mononucleosomes. In contrast, Jarid2 was not (Figure 4D). The enrichment of Ezh2, but not Jarid2 on H3.3K27M mono-nucleosomes was confirmed by Western blot (Figure S4). We noticed that another PRC2 core subunit, RbAp46/48, bound to wild type and mutant mononucleosome similarly, which is likely due to the fact that these two proteins are present in other chromatin complexes including CAF-1 (Verreault et al., 1996) and histone deacetylase complex NuRD (Xue et al., 1998; Zhang et al., 1999). Together, these results support the idea that Jarid2 can relieve the inhibitory effect of H3.3K27M mutant proteins on PRC2 core complex.

### Targeting Jarid2 to the H3.3K27M enriched loci leads to increased Ezh2 and H3K27me3 locally

To directly test the idea that Jarid2 can help overcome the H3.3K27M-mediated inhibition of PRC2 in cells, we used the CRISPR/dCas9 targeting system (Gilbert et al., 2013) to target Ezh2 or Jarid2 to two loci enriched with H3.3K27M, but with low levels of H3K27me3 in SF8628. The dCas9 (dCas9-Flag), which lacks nuclease activity, was fused to Ezh2 (dCas9-Ezh2-Flag) or Jarid2 (dCas9-Jarid2-Flag) (Figure S5A). Five sgRNAs surrounding the *PLXNA2* promoter were tested and one sgRNA with the highest targeting efficiency was selected for further experiments (Figure S5B). After transfection, the enrichments of Jarid2, Ezh2 and H3K27me3 at regions flanking the targeting site were analyzed by ChIP-PCR. Compared to dCas9-Flag, dCas9-Ezh2-Flag and dCas9-Jarid2-Flag appeared to show broad peaks surrounding the targeting site, suggesting that dCas9-Ezh2-Flag and dCas9-Jarid2-Flag can spread once targeted to this locus using CRISPR/dCas9 (Figure 5A, upper panel). Targeting Jarid2 to the *PLXNA2* promoter resulted in an increase in both Ezh2 and H3K27me3 around the targeting loci (Figure 5A, middle two panels), while having no apparent effect on the levels of H3.3K27M mutant proteins at the *PLXNA2* promoter (Figure 5A, bottom panel). In contrast, expression of dCas9-Ezh2-Flag fusion protein or dCas9-Flag proteins alone without sgRNA did not affect the levels of Jarid2 or H2K27me3 at the *PLXNA2* promoter (Figure 5A, S5C-S5F). Consistent with increased H3K27me3, targeting Jarid2, but not Ezh2, by CRISPR-dCas9 to the *PLXNA2* promoter led to reduced expression of *PLXNA2* compared to targeting dCas9 itself (Figure 5B). These results strongly support the idea that Jarid2 can recruit Ezh2 to regions with H3.3K27M and relieve the inhibitory effect of the mutant proteins.

**Figure 5.**
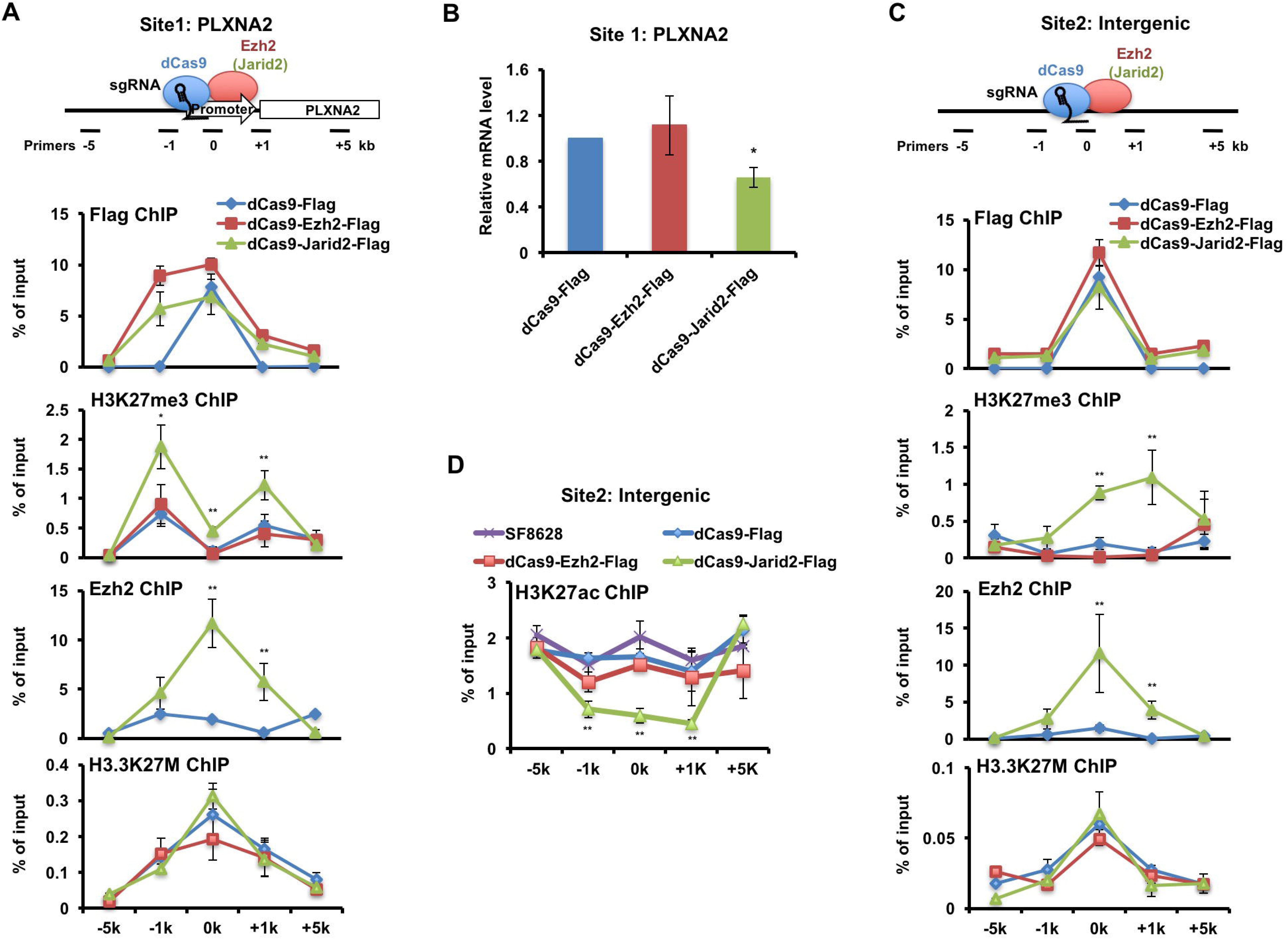
Targeting Jarid2 to H3.3K27M enriched loci results in localized increase in Ezh2 and H3K27me3. (**A**) Top: a schematic strategy of targeting Jarid2 or Ezh2 using CRISPR/dCas9 to the *PLXNA2* promoter enriched with H3.3K27M mutant proteins and with low levels of H3K27me3. The location of PCR primers used for analysis of ChIP DNA was also shown, with the CRISPR targeting site set as 0 kb. Bottom panels: ChIP-PCR analysis of dCas9 and its fusion proteins (dCas9-Ezh2-Flag and dCas9-Jarid2-Flag; Flag ChIP, upper panel), H3K27me3 (H3K27me3 ChIP, second panel), Ezh2 (Ezh2 ChIP) and H3.3K27M (H3K27M ChIP) at sites surrounding the CRISPR targeting sites in SF8628 cells expressing with dCas9, dCas9-Ezh2-Flag or dCas9-Jarid2-Flag. Data are mean ± SD (N = 3 independent replicates, ^*^P < 0.05, ^**^P < 0.01). (**B**) The expression of *PLXNA2* is reduced by targeting dCas9-Jarid2-Flag to the *PLXNA2* promoter. The expression of *PLXNA2* was analyzed by quantitative RT-PCR. dCas9-Flag, dCas9-Ezh2-Flag and dCas9-Jarid2-Flag were targeted to the PLXNA2 promoter as in (A). Data are mean ± SD (N = 3 independent replicates, ^*^P < 0.05). (**C**) Targeting Jarid2 to the intergenic region with H3.3K27M also results in increased H3K27me3. Top panel: the location of PCR primers at the intergenic region used for analysis of ChIP DNA was shown. The experiments were performed as described in A. Data are mean ± SD (N = 3 independent replicates, ^*^P < 0.05, ^**^P < 0.01). (**D**) H3K27ac is reduced by targeting Jarid2 to the intergenic region. H3K27ac ChIP was performed in SF8628 cells or cells expressing dCas9-Flag, dCas9-Ezh2-Flag or dCas9-Jarid2-Flag and analyzed using PCR primers outlined in C. Data are mean ± SD (N = 3 independent replicates, ^**^P < 0.01).

Similar results were observed when targeting Jarid2 to an intergenic region associated with H3.3K27M mutant proteins. Targeting Jarid2 but not Ezh2 by CRISPR/dCas9 resulted in an increase of H3K27me3 and a reduction of H3K27ac surrounding the targeting site despite the fact that Ezh2 levels flanking the targeted region were detected in both cases (Figure 5C-D). Control experiments that did not express sgRNAs did not show enrichment of Jarid2 or Ezh2 at the intergenic region nor changes in H3K27me3, Ezh2 and Jarid2 (Figure S5C-S5E and S5G). Together, these results demonstrate that targeting Jarid2 can recruit Ezh2 to H3.3K27M enriched promoter and intergenic region, which leads to an increase in H3K27me3 at these sites.

### Many putative tumor suppressor genes are silenced in DIPG cells through the H3K27me3-mediated mechanism

H3K27me3 is associated with gene silencing (Barski et al., 2007; Cao et al., 2002). We hypothesize that the presence of H3K27me3 in DIPG cells represses the expression of tumor suppressor genes (TSGs). To test this idea, we first asked whether the 676 genes with H3K27me3 peaks in three samples(SF8628, SF7761 and the primary DIPG tissue) we analyzed are putative TSGs. Based on the annotation of three different databases, we found that 80 genes including several noncoding RNAs were classified as TSGs (Figure S6A). To identify the candidate TSGs whose silencing is important for the proliferation of DIPG cells, we focused on analysis of 12 putative TSGs including well-known TSGs p15 and p16 (Figure S6A) that showed low expression in SF7761 and SF8628 cells based on RNA-seq. First, we tested their expression levels in 5 DIPG lines (SF8628, SF7761, DIPG17, DIPG13, and PED8) with H3.3K27M mutation, one line with H3.1K27M mutation (DIPG4), one line with H3.3G34V mutation (KNS42) and one brain tumor line with wild type H3.3 (SF9427). While the expression of these 12 genes was low in general in all these lines, clustering analysis showed that the expressions of NGFR, WT1 and MME were lowest among these 12 genes in these tumor cells, irrespectively mutation status of histone H3 when compared with human NSCs (Figure 6A). Next, we analyzed the enrichment of H3K27me3, Ezh2, and Jarid2 at the promoters of NGFR, WT1 and MME in these cell lines and human NSC and observed that Jarid2, H3K27me3 and Ezh2 were enriched at the promoters of WT1 promoter more than other two gene promoters (Figure S6B-D).

**Figure 6.**
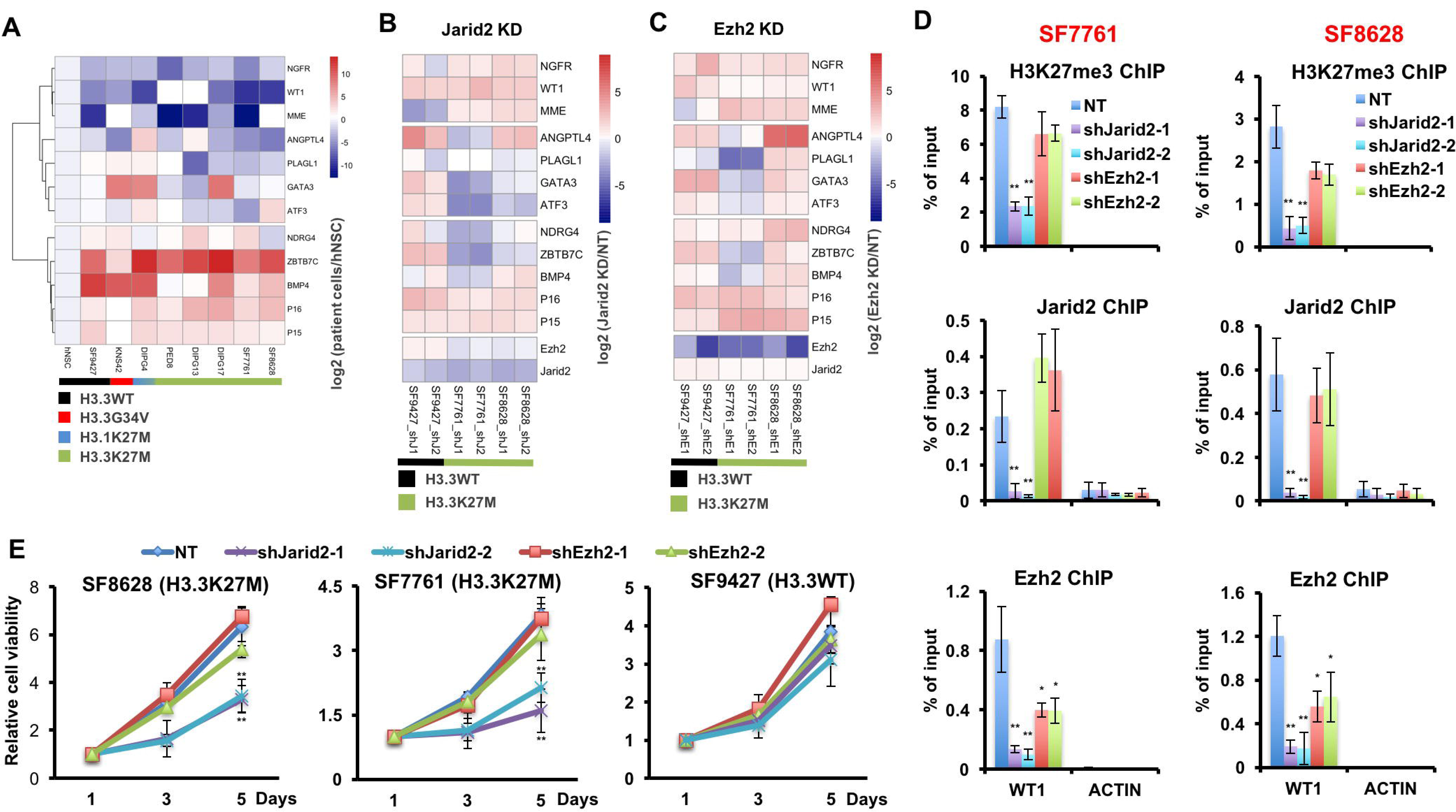
TSGs are silenced through H3K27me3-mediated mechanism in DIPG cells with H3.3K27M mutation. (**A**) The expression levels of 12 different tumor suppressor genes H3K27me3 at their promoters based on ChIP-seq in different DIPG lines and human NSC. Expressions of 12 different tumor suppressor genes were analyzed by quantitative RT-PCR and clustered hierarchically. Data are mean (N = 3 independent replicates). White blocks represent the ones whose gene expression was not detected. **(B and C**) Depletion of Jarid2 and Ezh2 has differential effects on the expression of WT1 in DIPG lines. The expression levels of different TSGs were analyzed in H3.3WT tumor cells (SF9427) or H3.3K27M mutant tumor cells (SF7761 and SF8628) after depletion of Ezh2 (B) and Jarid2 (C). Data are mean (N = 3 independent replicates). (**D**) Depletion of Jarid2 but not Ezh2 leads to reduction of H3K27me3 at the WT1 promoter in both SF7761 (left) and SF8682 (right) cells. Ezh2 and Jarid2 were depleted in SF7761 and SF8628 cells by two independent shRNAs. The H3K27me3, Jarid2 and Ezh2 levels at the promoter of WT1 were analyzed by ChIP-PCR. The enrichment of different proteins at the ACTIN promoter was used as a negative control. Data are mean ± SD (N = 3 independent replicates, ^*^P < 0.05, ^**^P < 0.01). NT, Non-Targeted control. (**E**) Depletion of Jarid2 reduces the proliferation of SF7761 and SF8628 cells. The proliferation of SF7761 and SF8628 cells was analyzed after Ezh2 or Jarid2 was depleted using two different shRNAs. NT: Non-target (NT) control. Data are mean ± SD (N = 3 independent replicates, ^**^P < 0.01).

Next, we evaluated the effect of depletions of Jarid2 and Ezh2 on the expression of these 12 genes in two DIPG lines (SF7761 and SF8628) and one tumor line with wild type H3.3 (SF9427). Depletion of Jarid2 using two independent shRNAs resulted in increased expression of p15, p16, WT1 and NGFR in all these tumor lines tested. These results indicate that Jarid2 is involved in silencing of several TSGs in DIPG cells (Figure 6B). Depletion of Ezh2 also resulted in increased expression of p15, p16 and NGFR in all tested tumor cells, whereas the expression of WT1 was not affected significantly after Ezh2 depletion (Figure 6C). ChIP-PCR analysis showed that depletion of Jarid2 resulted in reduced H3K27me3 and Ezh2 at the WT1 promoters, whereas depletion of Ezh2 had a less effect on H3K27me3 and Ezh2 levels at the promoter of WT1 despite the fact that depletion of Ezh2 led to reduced Ezh2 and H3K27me3 as determined by Western blot (Figure 6D and Figure 3A). These results indicate that Jarid2 and Ezh2 are involved in silencing of multiple TSGs in DIPG cells and that depletion of Jarid2 and Ezh2 have distinct effects on H3K27me3 levels at the WT1 promoter.

### Depletion of Jarid2 and Ezh2 has a differential effect on cell growth

Next, we analyzed how depletion of Jarid2 and Ezh2 affected the proliferation of two H3.3K27M mutant cells (Figure 6E). We observed that the growth of SF7761 and SF8628 was inhibited by depletion of Jarid2. Depletion of Ezh2 by two independent shRNAs had minor effects in this short-term culture (Mohammad et al., 2017; Piunti et al., 2017). In addition, depletion of Jarid2 and Ezh2 had no apparent effect on the growth of H3.3 wild type cells SF9427, despite the fact that the expressions of TSGs were activated in SF9427 cell line, suggesting that Jarid2 and Ezh2 mediated silencing of these TSGs is less critical for the proliferation of SF9427 line than two DIPG lines with the H3.3K27M mutation.

### Forced expression of WT1 inhibits the proliferation of H3.3K27M mutant cells

WT1 was first discovered to be mutated in Wilms Tumor, a pediatric kidney tumor (Gessler et al., 1990; Huff, 1998; Maiti et al., 2000). Since depletion of Jarid2 and Ezh2 had different effects on the expression of WT1 as well as proliferation, we asked whether forced expression of WT1 affected the proliferation of DIPG cells in a manner similar to Jarid2 depletion. We used two methods to increase WT1 expression. First, we overexpressed each of the four different isoforms of WT1 in H3.3K27M mutant cells (SF7761 and SF8628) and H3.3WT cells (SF9427), and found that the expression of each WT1 isoform inhibited the proliferation of two H3.3K27M mutant cells, but not H3.3WT cells (SF9427) (Figure 7A and 7B), suggesting that silencing WT1 is more important for the proliferation of DIPG cells. Second, we tested whether alterations in chromatin states of the WT1 promoter by targeting p300 using CRISPR/dCas9 would also change the expression of WT1 and cell proliferation. We targeted dCas9-HA and dCas9-HA fused with p300 catalytic domain (dCas9-HA-p300-HA) to the WT1 promoter in SF8628 cells using two independent sgRNAs (Figure S7A and S7B). The dCas9-HA and dCas9-HA-p300-HA were enriched at the WT1 promoter only when they were co-expressed with either of the two sgRNAs, but not at the promoter of ACTIN (Figure 7C, top panel). Targeting dCas9-HA-p300-HA to the WT1 promoter led to a reduction of H3K27me3, Ezh2 and Jarid2 as well as an increase in H3K27ac at the WT1 promoter, while having no apparent effect on the H3K27ac status of ACTIN promoter (Figure 7C). Moreover, the expression levels of WT1 increased through targeting dCas9-HA-p300-HA to the WT1 promoter (Figure 7D). Finally, the proliferation of SF8628 cells was inhibited when p300 catalytic domain was targeted to the WT1 promoter using CRISPR/dCas9 (Figure 7E). Together, these results show that alterations of chromatin states of WT1 promoter can change the expression of WT1 and proliferation of DIPG cells, supporting the idea that H3K27me3-mediated silencing of WT1 supports the proliferation of DIPG cells.

**Figure 7.**
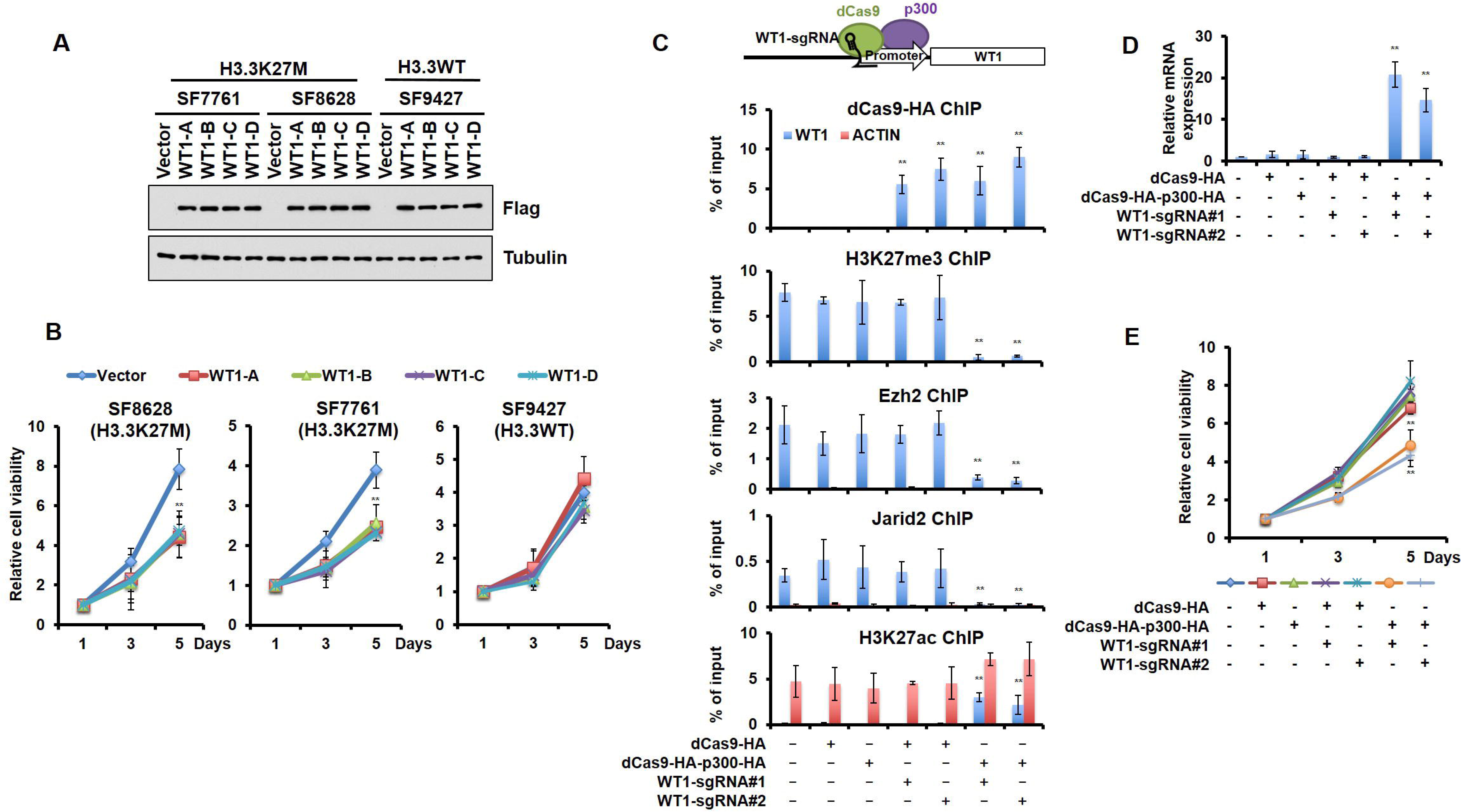
An increase in WT1 expression inhibits the proliferation of H3.3K27M mutant cells. (**A-B**) Cell proliferation of two DIPG lines were inhibited by overexpression of different WT1 isoforms. The expression of different Flag-tagged WT1 isoforms in two DIPG lines (SF7761 and SF8628) and one tumor line with wild type H3.3 (SF9427) was analyzed by Western blot using the indicated antibodies (A). Cell proliferation at different days of expression was monitored by cell titer-blue assay (B). Data are mean ± SD (N = 3 independent replicates, ^**^P < 0.01). (**C**) Targeting p300 to the WT1 promoter results an increase in H3K27ac and a reduction of H3K27me3. Top: a schematic strategy for targeting dCas9-HA-p300-HA fusion proteins to the promoter of WT1. ChIP-PCR analysis of dCas9-HA and dCas9-HA-p300-HA fusion proteins, H3K27me3, Ezh2, Jarid2 and H3K27ac at the promoter of WT1 was performed using cells transfected with different combinations of dCas9-HA, dCas9-HA-p300-HA and two different sgRNAs targeting the WT1 promoter shown in the bottom. Data are mean ± SD (N = 3 independent replicates, ^**^P < 0.01). (**D**) The expression of WT1 increases after targeting dCas9-HA-p300-HA to the WT1 promoter. The expression of WT1 was analyzed by quantitative RT-PCR. Data are mean ± SD (N = 3 independent replicates, ^**^P < 0.01). (**E**) SF8628 cell proliferation is inhibited by targeting dCas9-HA-p300-HA to the WT1 promoter. Data are mean ± SD (N = 3 independent replicates, ^**^P < 0.01).

## Discussion

How local H3K27 methylation is present in an environment of global loss of H3K27 methylation in cells with the H3K27M mutation is largely unknown. Here we show that the levels of H3.3K27M mutant proteins at gene promoters with H3K27me3 are low. Furthermore, we show that Jarid2, a partner of the PRC2 complex, can relieve the inhibition of PRC2 by H3.3K27M mutant proteins *in vitro* and *in vivo.* Finally, we show that many TSGs are silenced in DIPG cells through an H3K27 methylation and Jarid2 mediated mechanism. Forced expression of WT1, one of TSGs silenced in DIPG lines inhibits proliferation of DIPG cells. Together, these studies provide the mechanistic insight into the local retention of H3K27me3 in the environment of the global reduction and the significance of retaining H3K27me3 in DIPG tumor cells

### Two mechanisms for the local retention of H3K27me3 in DIPG cells

Since the identification of H3.3K27M mutation in DIPG cells, we and others have shown that H3.3K27M mutant proteins drive the global loss of H3K27me3 on wild type histone proteins in part through its inhibitory effect on the enzymatic activity of the PRC2 complex (Bender et al., 2013; Chan et al., 2013b; Lewis et al., 2013). In the environment of global loss of H3K27me3 in DIPG tumors, we and others also observed that H3K27me3 peaks were present at hundreds of loci based on analysis of H3K27me3 ChIP-seq (Bender et al., 2013; Chan et al., 2013b). In this study, we further confirm the dichotomous changes of H3K27me3 in one additional DIPG xenograft line and one primary tumor. Importantly, we show that a significant number of gene promoters with H3K27me3 are common among three DIPG samples analyzed (Figure 1B), suggesting that the retention of H3K27me3 at these loci likely have a role in DIPG tumor cells.

We have provided multiple lines of evidence supporting the idea that there are at least two mechanisms by which H3K27me3 is present at hundreds of loci in the environment of global loss. First, the levels of H3.3K27M mutant proteins at these loci with H3K27me3 are low compared to those without H3K27me3. The structure of PRC2 and H3.3K27M peptides reveals that methionine 27 is well accommodated by the hydrophobic side chains at the Ezh2 active site (Jiao and Liu, 2015; Justin et al., 2016). The energy cost of de-solvating methionine from the active site is much higher than lysine, which explains the higher binding affinity of PRC2 with H3.3K27M mutant protein or peptide than the corresponding wild type one. Thus, H3.3K27M mutant proteins likely function as competitive inhibitors of the PRC2 enzyme. Consistent with this idea, replacement of other histone lysine residues to methionine such as H3 lysine 9 and histone H3 lysine 36 also inhibits enzymatic activity of their corresponding lysine methyltransferases (Fang et al., 2016; Jayaram et al., 2016; Lu et al., 2016; Shan et al., 2016). In fission yeast, overexpression of Clr4, the H3K9 methyltransferase, overcomes the inhibition of H3K9M mutant proteins on H3K9me3 (Shan et al., 2016). Thus, levels of H3.3K27M mutant proteins detected at most gene promoters with H3K27me3 may be insufficient to inhibit the enzymatic activity of PRC2 complex at these gene promoters.

Why then are the H3.3K27M mutant proteins low at these loci? H3.3 is assembled into nucleosomes in a replication-independent pathway (Ahmad and Henikoff, 2002b). H3.3 ChlP-seq in different cell lines reveals that H3.3 is enriched at actively transcribed genes compared to lowly expressed genes (Chow et al., 2005; McKittrick et al., 2004; Schübeler et al., 2004;Stroud et al., 2012). We have shown that H3.3K27M mutant proteins exhibit a similar pattern of localization in two DIPG lines, suggesting that H3.3K27M mutant proteins are assembled into nucleosomes in DIPG cells like WT H3.3. Therefore, the low levels of H3.3K27M mutant proteins at genes with H3K27me3 likely reflect the fact that the expression of these genes is low in tumor initiating cells for DIPG tumors.

In addition to the low levels of H3.3K27M mutant proteins, we also presented several lines of evidence supporting the idea that Jarid2, when formed a complex with the core PRC2 subunits, can target Ezh2 to the loci close to H3.3K27M mutant proteins and relieve the inhibitory effect of H3.3K27M mutant proteins. First, we show that at similar levels of H3.3K27M mutant proteins, H3K27me3 levels are high at promoter regions with Jarid2 compared to those without Jarid2 in DIPG cells. Second, H3.3K27M peptide inhibits the enzymatic activity of Jarid2-PRC2 complex less efficiently than PRC2 alone *in vitro.* Third, targeting Jarid2 to two chromatin loci with H3.3K27M mutant proteins leads to increased recruitment of Ezh2 and increased enrichment of H3K27me3. These results strongly support the model that Jarid2 overcome the inhibition of PRC2 activity by H3.3K27M mutant proteins.

DIPG tumors occur primarily in mid-line brain structures and almost exclusively in children (Jones and Baker, 2014), suggesting these tumors arise from a specific lineage of neuronal precursor within these regions (Funato et al., 2014). We and others have shown that expression of H3.3K27M mutant proteins results in a global loss of H3K27 methylation, irrespective of cell-type or organisms (Chan et al., 2013b; Herz et al., 2014; Lewis et al., 2013). Therefore, the global reduction of H3K27 methylation in DIPG cells may not be the sole factor contributing to the tissue- and location-specific transformation observed in DIPG cells. Jarid2 can target PRC2 to specific chromatin loci and stimulate enzymatic activity of the PRC2 core complex. The expression of Jarid2 is high in mES cells and is low in differentiated cells (Li et al., 2010). Therefore, we propose that the tissue specific expression of Jarid2 can help locus-specific retention of H3K27 methylation and transform neuronal progenitor cells within mid-line regions into DIPG tumors.

### Epigenetic silencing of TSGs contributes to the development of DIPG cells

Is there any function of the retention of H3K27me3 in DIPG cells? Since H3K27me3 is involved in gene silencing, we hypothesize that retention of H3K27me3 in DIPG cells may help silence TSGs. Indeed, we observed that over 80 putative TSGs have high levels of H3K27me3 at their promoters in all three DIPG samples tested. It is known that inactivation of TSGs by genetic and epigenetic means is important for both tumor initiation and proliferation of tumor cells (Dawson and Kouzarides, 2012; Nephew and Huang, 2003; Riley and Anderson, 2011). In this study, we focus on identifying genes that when silenced are involved in proliferation of DIPG cells. We show that forced expression of WT1 either using a plasmid or alterations of chromatin state of WT1 endogenous promoter inhibit the proliferation of DIPG cells. These results strongly indicate that H3K27me3-mediated epigenetic silencing of WT1 is important for the proliferation of DIPG cells. WT1 is a zinc finger transcription factor (Bardeesy and Pelletier, 1998). It was first identified as a TSG in Wilms tumor, a tumor also found exclusively in children (Gessler et al., 1990; Maiti et al., 2000). Accumulating evidence suggests that WT1 is also expressed in many different classes of intracranial tumors, including gliomas (Izumoto et al., 2008), oligodendrogliomas (Rauscher et al., 2014), ependymomas (Yeung et al., 2013), and meningiomas (Iwami et al., 2013) and functions as an oncogene during tumor initiation and progression of these tumors. Therefore, it is interesting that WT1 functions as a TSG that modulates the proliferation of DIPG cells.

In addition to WT1, other TSGs including cell cycle inhibitors p16 and p15 also contain H3K27me3 at their promoters and are expressed at low levels in DIPG cells. Moreover, depletion of Jarid2 and Ezh2 also results in increased expression of 5 TSGs out of 12 genes tested. These results indicate that these genes are also silenced through the H3K27me3-mediated mechanism. We propose that H3K27me3-mediated silencing of TSGs likely contributes to the initiation and maintenance of DIPG tumors despite the fact that H3K27me3 is reduced globally. Consistent with this idea, inactive mutations at Ezh2, catalytic subunit of PRC2, have not reported despite the fact that many DIPG tumor samples have been sequenced. Therefore, we propose that it is the dichotomous changes in H3K27me3, including both global loss and locus-specific retention/gain of H3K27me3 in H3K27me3 in DIPG cells contribute to tumorigenesis.

The dichotomous change in H3K27me3 in DIPG cells is reminiscent of DNA methylation alteration observed in many cancer cells (Baylin and Jones, 2011). It is known for a long time that genomic DNA in most cancer cells is hypomethylated (Eden et al., 2003; Ehrlich, 2009), and yet DNA hypermethylation at TSGs has been frequently reported in many cancer types (Baylin, 2005; Esteller, 2007; Struhl, 2014). DNA methylation analysis in DIPG cells also indicates that DNA is hypomethylated (Bender et al., 2013). It would be interesting to determine whether DNA is hypermethylated at TSG genes or any genes with high levels of H3K27 methylation in the future.

In summary, we have shown that H3K27me3 is detected in two DIPG cells and one primary DIPG tissue, and a large number of gene promoters with H3K27me3 are the same among these three samples. The presence of H3K27me3 at these gene promoters is due to low levels of H3.3K27M mutant proteins and/or the formation of Jarid2-PRC2 complex, which can relieve the inhibitory effect of H3.3K27M mutant proteins. Finally, we show that H3K27me3-mediated silencing of WT1 is required for the proliferation of DIPG cells. These results indicate that inhibition of Ezh2 may also be an option for the treatment of DIPG tumors in the future.

## Author contributions

Conceptualization, Z.Z and D.F.; Methodology, D.F., and H.G.; Bioinformatic Analysis, H.G.; D.F. performed almost all experiments and L.C., J.L. and H.Z. performed some of the experiments. J.N.S. and D.J.D. provided critical resources. Figure S1A was performed in D.J.D. laboratory, D.F., H.G. and Z.Z wrote the original drafts. All authors read and J.N.S., D.J.D. and J.L. edited the manuscript.

## Acknowledgement

We thank Dr. Paul Grundy and Paul Goodyer for WT1 plasmids, and Ross Tomaino for SILAC mass spectrometry analysis. We thank Michelle Monje for DIPG cell lines. These studies were supported by the NIH CA157489 (Z.Z.). D.F. was supported by a fellowship from the Fraternal Order of Eagles Cancer Research Fund. ChIP-seq and RNA-seq data sets have been deposited into the Gene Expression Omnibus under accession number GSE94834.

## Reference

Adam, S., Polo, S., and Almouzni, G. 2013. Transcription recovery after DNA damage requires chromatin priming by the H3. 3 histone chaperone HIRA. Cell 155, 94–106.

Ahmad, K., and Henikoff, S. 2002a. Histone H3 variants specify modes of chromatin assembly. Proc. Natl. Acad. Sci. U. S. A 99 Suppl 4, 16477–16484.

Ahmad, K., and Henikoff, S. 2002b. The histone variant H3.3 marks active chromatin by replication-independent nucleosome assembly. Mol. Cell 9, 1191–1200.

Banaszynski, L.A., Wen, D., Dewell, S., Whitcomb, S.J., Lin, M., Diaz, N., Elsässer, S.J., Chapgier, A., Goldberg, A.D., Canaani, E., et al. 2013. Hira-Dependent Histone H3.3 Deposition Facilitates PRC2 Recruitment at Developmental Loci in ES Cells. Cell 155, 107–120.

Bardeesy, N., and Pelletier, J. 1998. Overlapping RNA and DNA binding domains of the wt1 tumor suppressor gene product. Nucleic Acids Res. 26, 1784–1792.

Barski, A., Cuddapah, S., Cui, K., Roh, T.Y., Schones, D.E., Wang, Z., Wei, G., Chepelev, I., and Zhao, K. 2007. High-Resolution Profiling of Histone Methylations in the Human Genome. Cell 129, 823–837.

Baylin, S.B. (2005). DNA methylation and gene silencing in cancer. Nat. Clin. Pract. Oncol. Oncol. 2 Suppl 1, S4–11.

Baylin, S.B., and Jones, P.A. 2011. A decade of exploring the cancer epigenome - biological and translational implications. Nat. Rev. Cancer 11, 726–734.

Bender, S., Tang, Y., Lindroth, A.M., Hovestadt, V., Jones, D.T.W., Kool, M., Zapatka, M., Northcott, P.A., Sturm, D., Wang, W., et al. 2013. Reduced H3K27me3 and DNA Hypomethylation Are Major Drivers of Gene Expression in K27M Mutant Pediatric High-Grade Gliomas. Cancer Cell 24, 660–672.

Burgess, R.J., and Zhang, Z. 2013. Histone chaperones in nucleosome assembly and human disease. Nat. Struct. Mol. Biol. 20, 14–22.

Calo, E., and Wysocka, J. 2013. Modification of Enhancer Chromatin: What, How, and Why? Mol. Cell 49, 825–837.

Campos, E.I., and Reinberg, D. 2009. Histones: annotating chromatin. Annu. Rev. Genet. 43, 559–599.

Cao, R., Wang, L., Wang, H., Xia, L., Erdjument-Bromage, H., Tempst, P., Jones, R.S., and Zhang, Y. 2002. Role of histone H3 lysine 27 methylation in Polycomb-group silencing. Science 298, 1039–1043.

Castel, D., Philippe, C., Calmon, R., Le Dret, L., Truffaux, N., Boddaert, N., Pagès, M., Taylor, K.R., Saulnier, P., Lacroix, L., et al. 2015. Histone H3F3A and HIST1H3B K27M mutations define two subgroups of diffuse intrinsic pontine gliomas with different prognosis and phenotypes. Acta Neuropathol. 130, 815–827.

Chan, K.M., Han, J., Fang, D., Gan, H., and Zhang, Z. 2013a. A lesson learned from the H3.3K27M mutation found in pediatric glioma A new approach to the study of the function of histone modifications in vivo? Cell Cycle 12, 2546–2552.

Chan, K.M., Fang, D., Gan, H., Hashizume, R., Yu, C., Schroeder, M., Gupta, N., Mueller, S., David James, C., Jenkins, R., et al. 2013b. The histone H3.3K27M mutation in pediatric glioma reprograms H3K27 methylation and gene expression. Genes Dev 27, 985–990.

Chen, P., Zhao, J., Wang, Y., Wang, M., Long, H., Liang, D., Huang, L., Wen, Z., Li, W., Li, X., et al. 2013. H3.3 actively marks enhancers and primes gene transcription via opening higher-ordered chromatin. Genes Dev. 27, 2109–2124.

Chow, C.-M., Georgiou, A., Szutorisz, H., Maia e Silva, A., Pombo, A., Barahona, I., Dargelos, E., Canzonetta, C., and Dillon, N. 2005. Variant histone H3.3 marks promoters of transcriptionally active genes during mammalian cell division. EMBO Rep. 6, 354–360.

Creyghton, M.P., Cheng, A.W., Welstead, G.G., Kooistra, T., Carey, B.W., Steine, E.J., Hanna, J., Lodato, M. a, Frampton, G.M., Sharp, P. a, et al. 2010. Histone H3K27ac separates active from poised enhancers and predicts developmental state. Proc. Natl. Acad. Sci. U. S. A. 107, 21931–21936.

Dawson, M.A., and Kouzarides, T. 2012. Cancer epigenetics: From mechanism to therapy. Cell 150, 12–27.

Deaton, A.M., G??mez-Rodr??guez, M., Mieczkowski, J., Tolstorukov, M.Y., Kundu, S., Sadreyev, R.I., Jansen, L.E.T., and Kingston, R.E. 2016. Enhancer regions show high histone H3.3 turnover that changes during differentiation. Elife 5.

Dellaretti, M., Reyns, N., Touzet, G., and et al. 2012. Diffuse brainstem glioma: prognostic factors. J. Neurosurg 117, 810–814.

Duarte, L.F., Young, A.R.J., Wang, Z., Wu, H.-A., Panda, T., Kou, Y., Kapoor, A., Hasson, D., Mills, N.R., Ma’ayan, A., et al. 2014. Histone H3.3 and its proteolytically processed form drive a cellular senescence programme. Nat. Commun. 5, 5210.

Eden, A., Gaudet, F., Waghmare, A., and Jaenisch, R. 2003. Chromosomal Instability and Tumors Promoted by DNA Hypomethylation. Science (80-.). 300, 2003.

Ehrlich, M. 2009. DNA hypomethylation in cancer cells. Epigenomics 1, 239–259.

Elsässer, S.J., Noh, K.-M., Diaz, N., Allis, C.D., and Banaszynski, L.A. 2015. Histone H3.3 is required for endogenous retroviral element silencing in embryonic stem cells. Nature 522, 240–244.

Esteller, M. (2007). Epigenetic gene silencing in cancer: The DNA hypermethylome. Hum. Mol. Genet. 16.

Fang, D., Gan, H., Lee, J.-H., Han, J., Wang, Z., Riester, S.M., Jin, L., Chen, J., Zhou, H., Wang, J., et al. 2016. The histone H3.3K36M mutation reprograms the epigenome of chondroblastomas. Science 352, 1344–1348.

Fontebasso, A.M., Papillon-Cavanagh, S., Schwartzentruber, J., Nikbakht, H., Gerges, N., Fiset, P.-O., Bechet, D., Faury, D., De Jay, N., Ramkissoon, L.A., et al. 2014. Recurrent somatic mutations in ACVR1 in pediatric midline high-grade astrocytoma. Nat. Genet. 46, 462–466.

Frey, A., Listovsky, T., Guilbaud, G., Sarkies, P., and Sale, J.E. 2014. Histone H3.3 is required to maintain replication fork progression after UV damage. Curr. Biol. 24, 2195–2201.

Funato, K., Major, T., Lewis, P.W., Allis, C.D., and Tabar, V. 2014. Use of human embryonic stem cells to model pediatric gliomas with H3.3K27M histone mutation. Science (80-.). 346, 1529–1533.

Gessler, M., Poustka, A., Cavenee, W., Neve, R.L., Orkin, S.H., and Bruns, G.A. 1990. Homozygous deletion in Wilms tumours of a zinc-finger gene identified by chromosome jumping. Nature 343, 774–778.

Gilbert, L. a, Larson, M.H., Morsut, L., Liu, Z., Gloria, A., Torres, S.E., Stern-ginossar, N., Brandman, O., Whitehead, H., Doudna, J. a, et al. 2013. CRISPR-Mediated Modular RNA-Guided Regualtion of Transcription in Eukaryotes. Cell 154, 442–451.

Guo, R., Zheng, L., Park, J.W., Lv, R., Chen, H., Jiao, F., Xu, W., Mu, S., Wen, H., Qiu, J., et al. 2014. BS69/ZMYND11 reads and connects histone H3.3 lysine 36 trimethylation-decorated chromatin to regulated pre-mRNA processing. Mol. Cell 56, 298–310.

Heintzman, N.D., Hon, G.C., Hawkins, R.D., Kheradpour, P., Stark, A., Harp, L.F., Ye, Z., Lee, L.K., Stuart, R.K., Ching, C.W., et al. 2009. Histone modifications at human enhancers reflect global cell-type-specific gene expression. Nature 459, 108–112.

Herz, H.-M., Morgan, M., Gao, X., Jackson, J., Rickels, R., Swanson, S.K., Florens, L., Washburn, M.P., Eissenberg, J.C., and Shilatifard, A. 2014. Histone H3 lysine-to-methionine mutants as a paradigm to study chromatin signaling. Science (80-.). 345, 1065–1070.

Huff, V. (1998). Wilms tumor genetics. In American Journal of Medical Genetics, pp. 260–267.

Iwami, K., Natsume, A., Ohno, M., Ikeda, H., Mineno, J., Nukaya, I., Okamoto, S., Fujiwara, H., Yasukawa, M., Shiku, H., et al. 2013. Adoptive transfer of genetically modified Wilms’ tumor 1-specific T cells in a novel malignant skull base meningioma model. Neuro. Oncol. 15, 747–758.

Izumoto, S., Tsuboi, A., Oka, Y., Suzuki, T., Hashiba, T., Kagawa, N., Hashimoto, N., Maruno, M., Elisseeva, O. a, Shirakata, T., et al. 2008. Phase II clinical trial of Wilms tumor 1 peptide vaccination for patients with recurrent glioblastoma multiforme. Neurosurg. 108, 963–971.

Jacob, Y., Bergamin, E., Donoghue, M.T. a, Mongeon, V., LeBlanc, C., Voigt, P., Underwood, C.J., Brunzelle, J.S., Michaels, S.D., Reinberg, D., et al. 2014. Selective methylation of histone H3 variant H3.1 regulates heterochromatin replication. Science 343, 1249–1253.

Jang, C.W., Shibata, Y., Starmer, J., Yee, D., and Magnuson, T. 2015. Histone H3.3 maintains genome integrity during mammalian development. Genes Dev. 29, 1377–1393.

Jayaram, H., Hoelper, D., Jain, S.U., Cantone, N., Lundgren, S.M., Poy, F., Allis, C.D., Cummings, R., Bellon, S., Lewis, P.W., et al. 2016. S-adenosyl methionine is necessary for inhibition of the methyltransferase G9a by the lysine 9 to methionine mutation on histone H3. Pnas 113, 6182–6187.

Jenuwein, T., and Allis, C.D. 2001. Translating the histone code. Science (80-.). 293, 1074–1080.

Jiao, L., and Liu, X. 2015. Structural basis of histone H3K27 trimethylation by an active polycomb repressive complex 2. Science (80-.). 350, 291.

Jones, C., and Baker, S.J. 2014. Unique genetic and epigenetic mechanisms driving paediatric diffuse high-grade glioma. Nat Rev Cancer.

Justin, N., Zhang, Y., Tarricone, C., Martin, S.R., Chen, S., Underwood, E., De Marco, V., Haire, L.F., Walker, P.A., Reinberg, D., et al. 2016. Structural basis of oncogenic histone H3K27M inhibition of human polycomb repressive complex 2. Nat. Comm 7, 11316.

Kaneko, S., Son, J., Shen, S.S., Reinberg, D., and Bonasio, R. 2013. PRC2 binds active promoters and contacts nascent RNAs in embryonic stem cells. Nat. Struct. & Mol. Biol 20, 1258–1264.

Kaneko, S., Bonasio, R., Saldaña-Meyer, R., Yoshida, T., Son, J., Nishino, K., Umezawa, A., and Reinberg, D. 2014. Interactions between JARID2 and Noncoding RNAs Regulate PRC2 Recruitment to Chromatin. Mol. Cell 53, 290–300.

Kanhere, A., Viiri, K., Araújo, C.C., Rasaiyaah, J., Bouwman, R.D., Whyte, W.A., Pereira, C.F., Brookes, E., Walker, K., Bell, G.W., et al. 2010. Short RNAs Are Transcribed from Repressed Polycomb Target Genes and Interact with Polycomb Repressive Complex-2. Mol. Cell 38, 675–688.

Khalil, A.M., Guttman, M., Huarte, M., Garber, M., Raj, A., Rivea Morales, D., Thomas, K., Presser, A., Bernstein, B.E., van Oudenaarden, A., et al. 2009. Many human large intergenic noncoding RNAs associate with chromatin-modifying complexes and affect gene expression. Proc. Natl. Acad. Sci. U. S. A. 106, 11667–11672.

Klose, R.J., and Zhang, Y. 2007. Regulation of histone methylation by demethylimination and demethylation. Nat. Rev. Mol. Cell Biol 8, 307–318.

Lee, J.-S., and Zhang, Z. 2016. O-linked N-acetylglucosamine transferase (OGT) interacts with the histone chaperone HIRA complex and regulates nucleosome assembly and cellular senescence. Proc. Natl. Acad. Sci. U. S. A. 1600509113–.

Lewis, P.W., Müller, M.M., Koletsky, M.S., Cordero, F., Lin, S., Banaszynski, L.A., Garcia, B.A., Muir, T.W., Becher, O.J., and Allis, C.D. 2013. Inhibition of PRC2 activity by a gain-of-function H3 mutation found in pediatric glioblastoma. Science 340, 857–861.

Li, G., Margueron, R., Ku, M., Chambon, P., Bernstein, B.E., and Reinberg, D. 2010. Jarid2 and PRC2, partners in regulating gene expression. Genes Dev. 24, 368–380.

Lin, C.-J., Conti, M., and Ramalho-Santos, M. 2013. Histone variant H3.3 maintains a decondensed chromatin state essential for mouse preimplantation development. Development 140, 3624–3634.

Louis, D.N., Ohgaki, H., Wiestler, O.D., Cavenee, W.K., Burger, P.C., Jouvet, A., Scheithauer, B.W., and Kleihues, P. 2007. The 2007 WHO classification of tumours of the central nervous system. Acta Neuropathol. 114, 97–109.

Lu, C., Jain, S.U., Hoelper, D., Bechet, D., Molden, R.C., Ran, L., Murphy, D., Venneti, S., Hameed, M., Pawel, B.R., et al. 2016. Histone H3K36 mutations promote sarcomagenesis through altered histone methylation landscape. Science (80-.). 352, 844–849.

Maiti, S., Alam, R., Amos, C.I., and Huff, V. 2000. Frequent association of??-catenin and WT1 mutations in Wilms tumors. Cancer Res. 60, 6288–6292.

Margueron, R., and Reinberg, D. 2011. The Polycomb complex PRC2 and its mark in life. Nature 469, 343–349.

Maze, I., Noh, K.-M., Soshnev, A.A., and Allis, C.D. 2014. Every amino acid matters: essential contributions of histone variants to mammalian development and disease. Nat. Rev. Genet. 15, 259–271.

McKittrick, E., Gafken, P.R., Ahmad, K., and Henikoff, S. 2004. Histone H3.3 is enriched in covalent modifications associated with active chromatin. Proc. Natl. Acad. Sci. U. S. A. 101, 1525–1530.

Mendenhall, E.M., Koche, R.P., Truong, T., Zhou, V.W., Issac, B., Chi, A.S., Ku, M., and Bernstein, B.E. 2010. GC-rich sequence elements recruit PRC2 in mammalian ES cells. PLoS Genet 6, 1–10.

Mohammad, F., Weissmann, S., Leblanc, B., Pandey, D.P., Hojfeldt, J.W., Comet, I., Zheng, C., Johansen, J.V., Rapin, N., Porse, B.T., et al. (2017). EZH2 is a potential therapeutic target for H3K27M-mutant pediatric gliomas. Nat Med advance online publication.

Mysliwiec, M.R., Bresnick, E.H., and Lee, Y. 2011. Endothelial Jarid2/Jumonji is required for normal cardiac development and proper Notch1 expression. J. Biol. Chem. 286, 17193–17204.

Mysliwiec, M.R., Carlson, C.D., Tietjen, J., Hung, H., Ansari, A.Z., and Lee, Y. (2012). Jarid2 (Jumonji, AT rich interactive domain 2) regulates NOTCH1 expression via histone modification in the developing heart. J. Biol. Chem. 287, 1235–1241.

Nephew, K.P., and Huang, T.H.M. 2003. Epigenetic gene silencing in cancer initiation and progression. Cancer Lett. 190, 125–133.

Ng, R.K., and Gurdon, J.B. 2008. Epigenetic memory of an active gene state depends on histone H3.3 incorporation into chromatin in the absence of transcription. Nat. Cell Biol. 10, 102–109.

Pasini, D., Cloos, P.A.C., Walfridsson, J., Olsson, L., Bukowski, J.-P., Johansen, J. V., Bak, M., Tommerup, N., Rappsilber, J., and Helin, K. 2010. JARID2 regulates binding of the Polycomb repressive complex 2 to target genes in ES cells. Nature 464, 306–310.

Peng, J.C., Valouev, A., Swigut, T., Zhang, J., Zhao, Y., Sidow, A., and Wysocka, J. 2009. Jarid2/Jumonji Coordinates Control of PRC2 Enzymatic Activity and Target Gene Occupancy in Pluripotent Cells. Cell 139, 1290–1302.

Pirrotta, V. 1998. Polycombing the genome: PcG, trxG and chromatin silencing. Cell 93, 333–336.

Piunti, A., Hashizume, R., Morgan, M.A., Bartom, E.T., Horbinski, C.M., Marshall, S.A., Rendleman, E.J., Ma, Q., Takahashi, Y., Woodfin, A.R., et al. (2017). Therapeutic targeting of polycomb and BET bromodomain proteins in diffuse intrinsic pontine gliomas. Nat Med advance online publication.

Pokholok, D.K., Harbison, C.T., Levine, S., Cole, M., Hannett, N.M., Tong, I.L., Bell, G.W., Walker, K., Rolfe, P.A., Herbolsheimer, E., et al. 2005. Genome-wide map of nucleosome acetylation and methylation in yeast. Cell 122, 517–527.

Rai, T.S., Cole, J.J., Nelson, D.M., Dikovskaya, D., Faller, W.J., Vizioli, M.G., Hewitt, R.N., Anannya, O., McBryan, T., Manoharan, I., et al. 2014. HIRA orchestrates a dynamic chromatin landscape in senescence and is required for suppression of Neoplasia. Genes Dev. 28, 2712–2725.

Rauscher, J., Beschorner, R., Gierke, M., Bisdas, S., Braun, C., Ebner, F.H., and Schittenhelm, J. 2014. WT1 expression increases with malignancy and indicates unfavourable outcome in astrocytoma. J. Clin. Pathol. 67, 556–561.

Riley, L.B., and Anderson, D.W. (2011). Cancer epigenetics. In Handbook of Epigenetics, pp. 521–534.

Rocha, S.T. da, Edwards, C.A., Ito, M., Ogata, T., and Ferguson-Smith, A.C. 2008. Genomic imprinting at the mammalian Dlk1-Dio3 domain. Trends Genet. 24, 306–316.

Sanulli, S., Justin, N., Teissandier, A., Ancelin, K., Portoso, M., Caron, M., Michaud, A., Lombard, B., da Rocha, S.T., Offer, J., et al. 2015. Jarid2 Methylation via the PRC2 Complex Regulates H3K27me3 Deposition during Cell Differentiation. Mol. Cell 57, 769–783.

Schübeler, D., MacAlpine, D.M., Scalzo, D., Wirbelauer, C., Kooperberg, C., Van Leeuwen, F., Gottschling, D.E., O’Neill, L.P., Turner, B.M., Delrow, J., et al. 2004. The histone modification pattern of active genes revealed through genome-wide chromatin analysis of a higher eukaryote. Genes Dev. 18, 1263–1271.

Schwartzentruber, J., Korshunov, A., and Liu, X.Y. (2012). Driver mutations in histone H3. 3 and chromatin remodelling genes in paediatric glioblastoma. Nature.

Shan, C.M., Wang, J., Xu, K., Chen, H., Yue, J.X., Andrews, S., Moresco, J.J., Yates, J.R., Nagy, P.L., Tong, L., et al. 2016. A histone H3K9M mutation traps histone methyltransferase Clr4 to prevent heterochromatin spreading. Elife 5.

Shen, X., Kim, W., Fujiwara, Y., Simon, M.D., Liu, Y., Mysliwiec, M.R., Yuan, G.C., Lee, Y., and Orkin, S.H. 2009. Jumonji Modulates Polycomb Activity and Self-Renewal versus Differentiation of Stem Cells. Cell 139, 1303–1314.

Shen, Y., Yue, F., McCleary, D.F., Ye, Z., Edsall, L., Kuan, S., Wagner, U., Dixon, J., Lee, L., Lobanenkov, V. V, et al. 2012. A map of the cis-regulatory sequences in the mouse genome. Nature 488, 116–120.

Simon, J.A., and Kingston, R.E. 2013. Occupying Chromatin: Polycomb Mechanisms for Getting to Genomic Targets, Stopping Transcriptional Traffic, and Staying Put. Mol. Cell 49, 808–824.

Simon, J., Chiang, a, Bender, W., Shimell, M.J., and O’Connor, M. 1993. Elements of the Drosophila bithorax complex that mediate repression by Polycomb group products. Dev. Biol. 158, 131–144.

Solomon, D.A., Wood, M.D., Tihan, T., Bollen, A.W., Gupta, N., Phillips, J.J.J., and Perry, A. 2016. Diffuse Midline Gliomas with Histone H3-K27M Mutation: A Series of 47 Cases Assessing the Spectrum of Morphologic Variation and Associated Genetic Alterations. Brain Pathol. 26, 569–580.

Son, J., Shen, S.S., Margueron, R., and Reinberg, D. 2013. Nucleosome-binding activities within JARID2 and EZH1 regulate the function of PRC2 on chromatin. Genes Dev. 27, 2663–2677.

Stroud, H., Otero, S., Desvoyes, B., Ramírez-Parra, E., Jacobsen, S.E., and Gutierrez, C. 2012. Genome-wide analysis of histone H3.1 and H3.3 variants in Arabidopsis thaliana. Proc. Natl. Acad. Sci. U. S. A. 109, 5370–5375.

Struhl, K. (2014). Is DNA methylation of tumour suppressor genes epigenetic? Elife 3, e02475.

Sturm, D., Witt, H., Hovestadt, V., Khuong-Quang, D.A., Jones, D.T.W., Konermann, C., Pfaff, E., Tönjes, M., Sill, M., Bender, S., et al. 2012. Hotspot Mutations in H3F3A and IDH1 Define Distinct Epigenetic and Biological Subgroups of Glioblastoma. Cancer Cell 22, 425–437.

Suka, N., Suka, Y., Carmen, A.A., Wu, J., and Grunstein, M. 2001. Highly specific antibodies determine histone acetylation site usage in yeast heterochromatin and euchromatin. Mol. Cell 8, 473–479.

Szenker, E., Ray-Gallet, D., and Almouzni, G. 2011. The double face of the histone variant H3.3. Cell Res 21, 421–434.

Tagami, H., Ray-Gallet, D., Almouzni, G., and Nakatani, Y. 2004. Histone H3.1 and H3.3 Complexes Mediate Nucleosome Assembly Pathways Dependent or Independent of DNA Synthesis. Cell 116, 51–61.

Takeuchi, T., Yamazaki, Y., Katoh-Fukui, Y., Tsuchiya, R., Kondo, S., Motoyama, J., and Higashinakagawa, T. 1995. Gene trap capture of a novel mouse gene, jumonji, required for neural tube formation. Genes Dev 9, 1211–1222.

Takeuchi, T., Watanabe, Y., Takano-Shimizu, T., and Kondo, S. 2006. Roles of jumonji and jumonji family genes in chromatin regulation and development. Dev. Dyn 235, 2449–2459.

Tan, M., Luo, H., Lee, S., Jin, F., Yang, J.S., Montellier, E., Buchou, T., Cheng, Z., Rousseaux, S., Rajagopal, N., et al. 2011. Identification of 67 histone marks and histone lysine crotonylation as a new type of histone modification. Cell 146, 1016–1028.

Turner, B.M. 2000. Histone acetylation and an epigenetic code. BioEssays 22, 836–845.

Verreault, A., Kaufman, P.D., Kobayashi, R., and Stillman, B. 1996. Nucleosome Assembly by a Complex of CAF-1 and Acetylated Histones H3/H4. Cell 87, 95–104.

Wen, H., Li, Y., Xi, Y., Jiang, S., Stratton, S., Peng, D., Tanaka, K., Ren, Y., Xia, Z., Wu, J., et al. 2014. ZMYND11 links histone H3.3K36me3 to transcription elongation and tumour suppression. Nature 508, 263–268.

Woo, C.J., Kharchenko, P. V., Daheron, L., Park, P.J., and Kingston, R.E. 2010. A Region of the Human HOXD Cluster that Confers Polycomb-Group Responsiveness. Cell 140, 99–110.

Wu, G., Broniscer, A., McEachron, T.A., Lu, C., Paugh, B.S., Becksfort, J., Qu, C., Ding, L., Huether, R., Parker, M., et al. 2012. Somatic histone H3 alterations in pediatric diffuse intrinsic pontine gliomas and non-brainstem glioblastomas. Nat Genet 44, 251–253.

Wu, G., Diaz, A.K., Paugh, B.S., Rankin, S.L., Ju, B., Li, Y., Zhu, X., Qu, C., Chen, X., Zhang, J., et al. 2014. The genomic landscape of diffuse intrinsic pontine glioma and pediatric non-brainstem high-grade glioma. Nat. Genet. 46, 444–450.

Xue, Y., Wong, J., Moreno, G.T., Young, M.K., Côté, J., and Wang, W. 1998. NURD, a novel complex with both ATP-dependent chromatin-remodeling and histone deacetylase activities. Mol. Cell 2, 851–861.

Yeung, J.T., Hamilton, R.L., Okada, H., Jakacki, R.I., and Pollack, I.F. 2013. Increased expression of tumor-associated antigens in pediatric and adult ependymomas: Implication for vaccine therapy. J. Neurooncol 111, 103–111.

Zhang, R., Chen, W., and Adams, P.D. 2007. Molecular dissection of formation of senescence-associated heterochromatin foci. Mol. Cell. Biol. 27, 2343–2358.

Zhang, Y., Ng, H.H., Erdjument-Bromage, H., Tempst, P., Bird, A., and Reinberg, D. 1999. Analysis of the NuRD subunits reveals a histone deacetylase core complex and a connection with DNA methylation. Genes Dev. 13, 1924–1935.

Zhao, J., Ohsumi, T.K., Kung, J.T., Ogawa, Y., Grau, D.J., Sarma, K., Song, J.J., Kingston, R.E., Borowsky, M., and Lee, J.T. 2010. Genome-wide Identification of Polycomb-Associated RNAs by RIP-seq. Mol. Cell 40, 939–953.

